# A cocktail of SARS-CoV-2 spike stem helix domain and receptor binding domain human monoclonal antibodies prevent the emerge of viral escape mutants

**DOI:** 10.1101/2025.10.16.682699

**Authors:** Yao Ma, Chengjin Ye, Michael S. Piepenbrink, Sara H. Mahmoud, Anastasija Cupic, Esteban Castro, Nathaniel Jackson, Mahmoud Bayoumi, Alvaro S. Padron, Adolfo Garcia-Sastre, Gregory C. Ippolito, Mark R. Walter, James J. Kobie, Luis Martinez-Sobrido

## Abstract

Neutralizing antibodies (NAbs) targeting the spike (S) glycoprotein remain a crucial therapeutic strategy against severe acute respiratory syndrome coronavirus 2 (SARS-CoV-2). However, emerging viral variants have escaped all Food and Drug Administration (FDA)-approved NAb treatments, underscoring the urgent need for effective therapeutic alternatives. Using a nanoluciferase (Nluc)-expressing attenuated recombinant SARS-CoV-2 lacking the open reading frames (ORF) 3a and 7b (Δ3a7b-Nluc), we characterized resistance profiles of two broadly protective NAbs targeting the S receptor binding domain (RBD) in S1 (1301B7) and the stem helix domain (SH) in S2 (1249A8). Serial passaging of Δ3a7b-Nluc under selective pressure identified a 1301B7 antibody-resistant mutants (ARM-B7) harboring an RBD mutation (S371F) that conferred resistance to 1301B7 and other RBD-directed NAbs (Casirivimab, SC27 and Sotrovimab). In contrast, no ARM emerged under treatment with 1249A8, or an antibody cocktail of 1301B7 (RBD) + 1249A8 (SH). These findings demonstrate that S2 SH-targeting NAbs shows higher genetic barrier to resistance than S1 RBD-targeting NAbs, and that a NAbs cocktail therapy targeting the SARS-CoV-2 S1 RBD and S2 SH offers the most effective strategy to prevent the emergence of escape mutations. Together, our findings provide critical insights into developing next-generation resistance-evading NAb therapies against SARS-CoV-2, and potentially other coronaviruses, and demonstrate the value of using our attenuated viral platforms for the safe identification of ARM without the potential biosafety concerns of doing these experiments using wild-type (WT) forms of SARS-CoV-2.

**SIGNIFICANCE:** The clinical efficacy of early SARS-CoV-2 NAbs has been challenged by the emergence of escape viral variants, highlighting an urgent need to anticipate resistance. Using a luminescent attenuated SARS-CoV-2 platform, we profiled resistant mutations against two broadly protective SARS-CoV-2 NAbs. Passage of Δ3a7b-Nluc in the presence of a NAb targeting the RBD S1 domain (1301B7) readily selected for an ARM, whereas passage in the presence of an SH S2 domain NAb (1249A8) did not. Notably, a cocktail of 1301B7 and 1249A8 created a high barrier of selecting SARS-CoV-2 ARM, preventing the emergence of resistant variants. We identified an S371F mutation in the S1 RBD of ARM-B7 that confers resistance to 1301B7 and other S1 RBD-targeting NAbs. These results highlight the importance of combination therapies targeting both variable RBD S1 and conserved SH S2 domain for the efficient treatment of SARS-CoV-2 and to prevent the emerge of NAb-induced escape mutations.

## INTRODUCTION

Over the past five years, coronavirus disease 2019 (COVID-19) has been responsible of over 700 million infections and approximately 7 million deaths (1). Notably, severe acute respiratory syndrome coronavirus 2 (SARS-CoV-2), the virus responsible of the COVID-19 pandemic, continues to evolve (2–5). Vaccines developed against the original SARS-CoV-2 strain no longer protect against emerging viral variants (6–10). Similarly, monoclonal neutralizing antibodies (NAbs) granted emergency use authorization (EUA) in the United States (US) rapidly lost effectiveness against newly emerging SARS-CoV-2 variants, and clinical use authorization of NAbs was revoked in early 2023 (11–13). The emergence of SARS-CoV-2 mutant strains necessitated the continuous updating of vaccines. Therefore, how to maximize protection against the emergence of new variants and suppress the emergence of escape mutations induced after vaccination has become an issue for the treatment of SARS-CoV-2 infections.

The receptor-binding domain (RBD) of SARS-CoV-2 spike (S) glycoprotein S1 region is a critical target for NAbs due to its role in mediating viral entry via binding to the angiotensin-converting enzyme 2 (ACE2) receptor (14, 15). However, the accumulation of mutations in the RBD of S1 across emerging variants has rendered many NAb therapies ineffective (11–13). A human monoclonal NAb, 1301B7, was isolated from a convalescent individual following Omicron infection in spring of 2023, using RBD-ACE2 fusion protein-based B cell isolation to enrich for antibodies targeting conserved ACE2-binding epitopes (16). 1301B7 exhibits broad and potent neutralization against SARS-CoV-2 original WA.1 strain as well as multiple viral variants, including Omicron subvariants BA.5, XBB.1.5, and JN.1, due to its unique binding mechanism involving the VH1-69 heavy chain and a long CDRH3 loop that engages conserved RBD S1 residues while tolerating mutations at positions like 417 and 456 residues (16).

Compared to the highly variable S1, the S2 stem domain demonstrates significant evolutionary conservation across β-coronaviruses, positioning it as a critical target for universal coronavirus antibodies and vaccines development (17–21). 1249A8 is a human NAb targeting the conserved membrane-proximal S2 stem helix (SH) region, thereby circumventing the immune evasion mechanisms associated with the highly variable SARS-CoV-2 RBD S1, making it an important candidate for universal coronavirus therapeutics (22–24). By disrupting the secondary structure and refolding events required for coronavirus post-fusion S to initiate membrane fusion and ultimately infection, 1249A8 demonstrates broad and robust neutralizing activity against multiple β-coronaviruses, including SARS-CoV-2, SARS-CoV, and MERS-CoV (22, 23). For this reason, 1249A8 exhibits potent neutralizing activity against SARS-CoV-2 and SARS-CoV in different animal models, demonstrating its universal β-coronavirus therapeutic potential (23).

The Δ3a7b-Nluc platform is a luminescent, attenuated recombinant SARS-CoV-2 engineered to enable the safe and efficient identification of antiviral-resistant mutants while circumventing the biosafety risks associated with using wild-type (WT) virus (25, 26). This system combines two key features: First, deletion of open reading frame (ORF) 3a and 7b accessory proteins of SARS-CoV-2 USA-WA1/2020 strain (GenBank accession no. MN985325), which attenuates viral pathogenicity while preserving replication (25, 27, 28). Second, expression of Nluc for real-time, high-throughput easy quantification of viral infection dynamics (26). The system maintains susceptibility to clinically relevant Food and Drug Administration (FDA)-approved or investigational antiviral compounds while being attenuated, allowing drug resistance profiling without the biosafety concerns associated with using WT SARS-CoV-2.

In this study, we employed Δ3a7b-Nluc to evaluate the emergence of antibody resistance mutants (ARMs) through serial passaging under increasing concentrations of NAbs. The study focused on two broad NAbs: 1301B7 (targeting RBD S1) and 1249A8 (targeting SH S2) alone or in combination. Following seven rounds of selection of Δ3a7b-Nluc with 1301B7, we isolated an antibody-resistant (ARM-B7) exhibiting significantly reduced neutralization (NT_50_ increase > 360-fold). Next generation sequencing (NGS) identified a dominant non-synonymous RBD S1 mutation (S371F) in ARM-B7. In contrast, parallel passaging of Δ3a7b-Nluc in the presence of 1249A8 failed to yield viruses with significant resistance, suggesting a higher genetic barrier to escape neutralization by 1249A8. Additionally, serial passage of Δ3a7b-Nluc in the presence of both NAbs (1301B7+1249A8) avoided resistance development, without significant change in viral neutralization after seven viral passages. These findings demonstrate that S2 SH-targeting NAbs like 1249A8 exhibit superior resistance profiles compared to S1 RBD-targeting NAbs like 1301B7, and that a cocktail of S2 SH and S1 RBD targeting NAbs (1249A8 + 1301B7) can efficiently prevent the emergence of ARMs. Furthermore, we identified S371F mutation as an important mutation to reduce the neutralizing activity of S1 RBD-targeting NAbs. We also demonstrate that our attenuated recombinant SARS-CoV-2 expressing Nluc (Δ3a7b-Nluc) platform represents a safe option to identify ARMs from monoclonal NAbs, or potentially sera preparations, to avoid biosafety concerns of using WT SARS-CoV-2.

## RESULTS

### Conservation of antibody binding epitopes

NAbs 1301B7 and 1249A8 target distinct structural regions within the SARS-CoV-2 S protein (**Figure 1A**). 1301B7 binds a conformational epitope mainly made of 11 amino acids within the S1 RBD, while 1249A8 recognizes a linear epitope spanning 12 amino acid residues in the membrane-proximal S2 SH region (**Figure 1A**). Conservation analysis across major SARS-CoV-2 variants revealed that four residues within the 1301B7 epitope (positions 421, 453, 489, and 494) remain invariant, while the remaining seven amino acids exhibit variant-specific substitutions, with JN.1 accumulating the highest mutational burden (6 altered residues) (**Table 1**). In contrast, the epitope of 1249A8 shows complete conservation across all major SARS-CoV-2 variants, without observed mutations, highlighting its structural stability and potential as a resilient therapeutic target (**Table 1**). 1301B7 (16) and 1249A8 (23) individual neutralizing activity against SARS-CoV-2 was previously described and their potential synergistic neutralizing activity was evaluated using a checkerboard dilution assay (**Figure S1**). Serial dilutions of each NAb, expressed as multiples of its individual 50% neutralization titer (NT_50_), were combined and tested for their capacity to neutralize Δ3a7b-Nluc (**Figure S1**). The results demonstrate that the neutralization potency of 1301B7 was not compromised by co-incubation with 1249A8, and similarly, 1249A8 activity was independent of 1301B7 concentration. This absence of competitive inhibition implies that the two antibodies bind to independent epitopes. Notably, we observed a synergistic neutralizing activity when both NAbs were combined (**Figure S1**).

**Figure 1.**
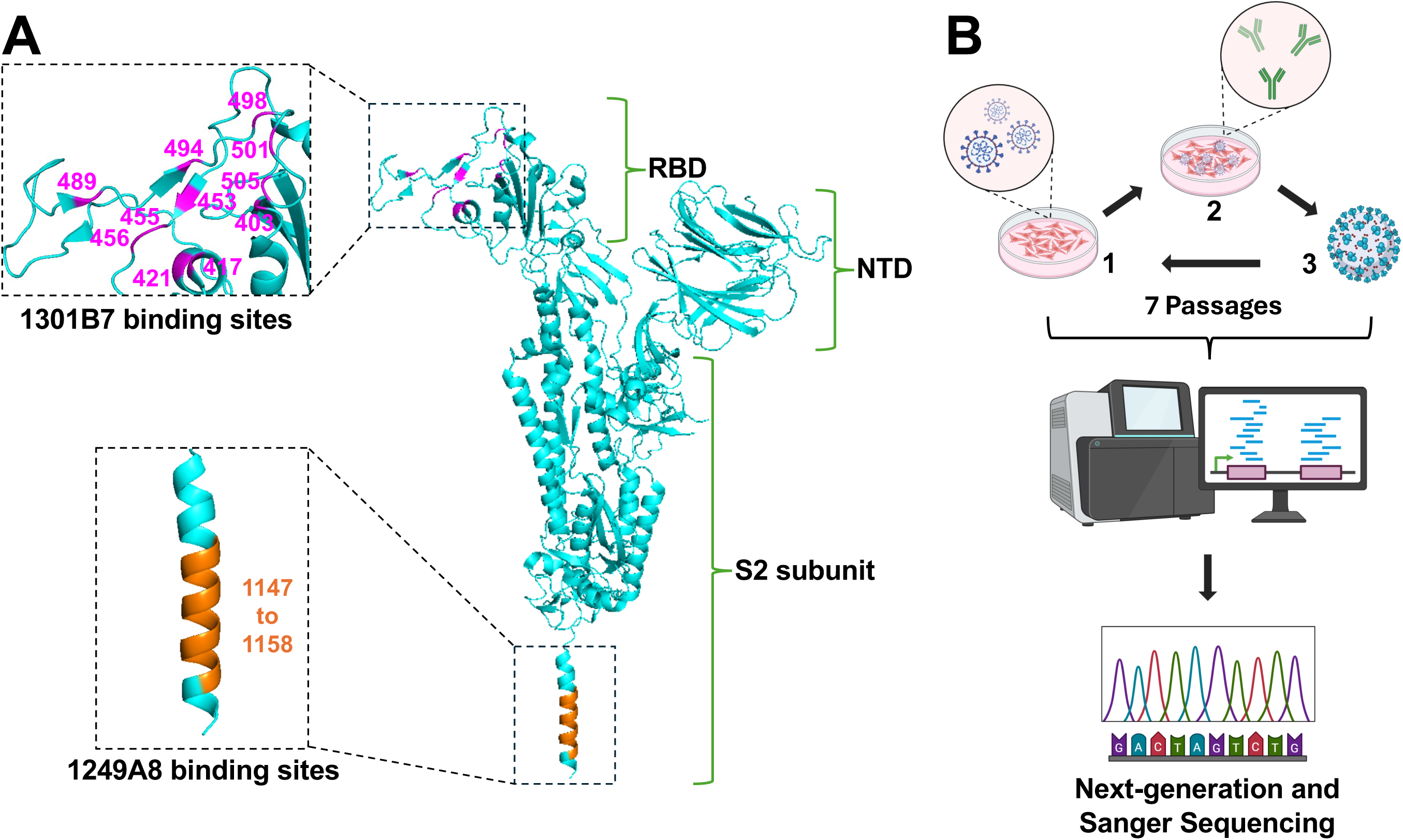
Neutralizing antibody (NAb) binding sites and selection of antibody-resistant mutants (ARMs). (A) Location of binding sites in SARS-CoV-2 S protein (cyan, PDBID:6XR8) for 1301B7 (magenta), which targets the S1 RBD, and 1249A8 (orange), which targets the S2 SH. (B) Schematic of ARM selection: The attenuated recombinant SARS-CoV-2 expressing nanoluciferase (Δ3a7b-Nluc) was serially passaged in Vero-AT cells under increasing concentrations of NAbs. RNA from the Passage 7 viral populations was subjected to next-generation sequencing (NGS) and Sanger sequencing to identify resistance-associated mutations.

**Table 1.** Conservation of 1301B7 and 1249A8 binding sites in SARS-CoV-2 variants.

|  | WA.1 | B.1.1.7 | B.1.351 | P.1 | AY.4 | BA.1 | BA.2 | BA.5 | EG.5 | XBB.1.5 | JN.1 |
| --- | --- | --- | --- | --- | --- | --- | --- | --- | --- | --- | --- |
| 1301B7 | R403 |  |  |  |  |  |  |  |  |  | R403K |
|  | K417 |  | K417N | K417T |  | K417N | K417N | K417N | K417N | K417N | K417N |
|  | Y421 |  |  |  |  |  |  |  |  |  |  |
|  | Y453 |  |  |  |  |  |  |  |  |  |  |
|  | L455 |  |  |  |  |  |  |  |  |  | L455S |
|  | F456 |  |  |  |  |  |  |  | F456L |  |  |
|  | Y489 |  |  |  |  |  |  |  |  |  |  |
|  | S494 |  |  |  |  |  |  |  |  |  |  |
|  | Q498 |  |  |  |  | Q498R | Q498R | Q498R | Q498R | Q498R | Q498R |
|  | N501 | N501Y | N501Y | N501Y |  | N501Y | N501Y | N501Y | N501Y | N501Y | N501Y |
|  | Y505 |  |  |  |  | Y505H | Y505H | Y505H | Y505H | Y505H | Y505H |
| 1249A8 | S1147 |  |  |  |  |  |  |  |  |  |  |
|  | F1148 |  |  |  |  |  |  |  |  |  |  |
|  | K1149 |  |  |  |  |  |  |  |  |  |  |
|  | E1150 |  |  |  |  |  |  |  |  |  |  |
|  | E1151 |  |  |  |  |  |  |  |  |  |  |
|  | L1152 |  |  |  |  |  |  |  |  |  |  |
|  | D1153 |  |  |  |  |  |  |  |  |  |  |
|  | K1154 |  |  |  |  |  |  |  |  |  |  |
|  | Y1155 |  |  |  |  |  |  |  |  |  |  |
|  | F1156 |  |  |  |  |  |  |  |  |  |  |
|  | K1157 |  |  |  |  |  |  |  |  |  |  |
|  | N1158 |  |  |  |  |  |  |  |  |  |  |

The information comes from outbreak.info (available online: https://outbreak.info/).

### Isolation of ARMs

To investigate the development of antibody resistance, Δ3a7b-Nluc was serially passaged in Vero-AT cells under increasing concentrations of 1301B7, 1249A8, or a cocktail of 1301B7 + 1249A8 (**Figures 1B and 2A**). Following seven serial passages, a variant with significant resistance to 1301B7 (ARM-B7) was isolated (**Figure 2B**). In contrast, parallel serial passage of Δ3a7b-Nluc under increasing concentrations of 1249A8 did not yield viruses with significant enhanced resistance (ARM-A8), indicating a higher genetic barrier to escape for this S2 SH region targeted by 1249A8 (**Figure 2B**). Notably, combination therapy with both NAbs effectively suppressed the emergence of resistance viruses (ARM-B7+A8), with no significant reduction in neutralization sensitivity observed after seven passages compared to the original Δ3a7b-Nluc (**Figure 2B**).

**Figure 2.**
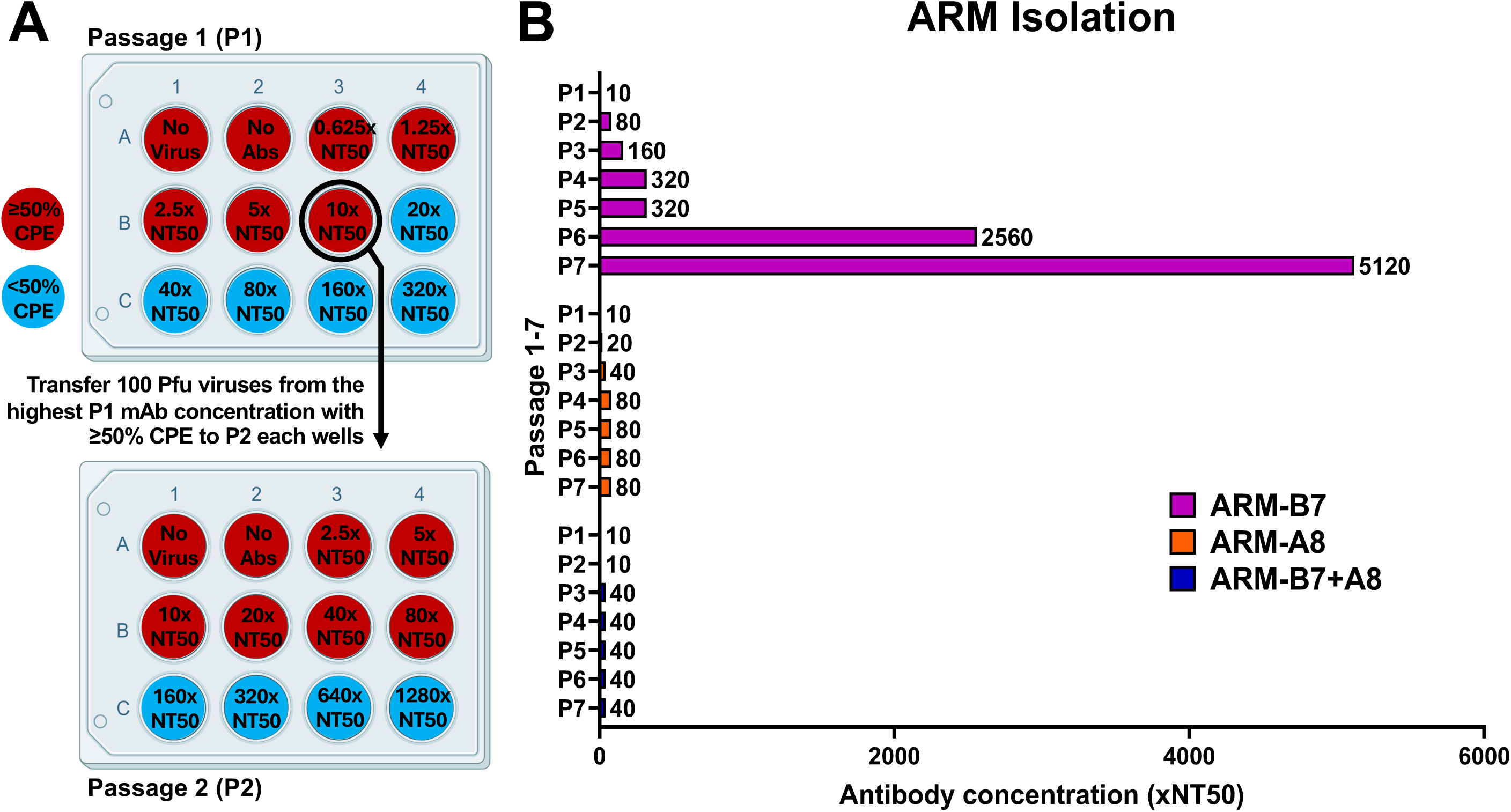
Isolation of ARMs. (A) Vero-AT cell monolayers were infected with Δ3a7b-Nluc (100–200 PFU/well) and treated with escalating concentrations of 1301B7 (NT_₅₀_ = 5.66 ng/mL) and 1249A8 (NT_₅₀_ = 2.09 µg/mL) alone or their combination. After 72 hours infection, cell culture supernatants from wells showing ≥50% cytopathic effect (CPE) at the highest NAb concentration were harvested to initiate the next passage. Viral load was quantified by Nluc activity. (B) Passaging result of ARMs isolation is shown as multiples of the antibody NT_50_ concentration.

To quantitatively assess the neutralization resistance of the ARMs, we performed plaque reduction neutralization tests (PRNT) to determine the NT_50_. ARM-B7 mutant exhibited a profound (∼360-fold increase) resistance to 1301B7 (NT_50_ = 2.04 µg/mL) compared to the parental Δ3a7b-Nluc virus (NT_50_ = 5.66 ng/mL) (**Figures 3A and 3C**). In contrast, ARM-A8 (NT_50_ = 7.93 ng/mL) and ARM-B7+A8 (NT_50_ = 8.17 ng/mL) mutants remain similar neutralized by 1301B7, with less than 2-fold increase in NT_50_ (**Figures 3A and 3C**). Importantly, ARM-B7, ARM-A8 and ARM-B7+A8 mutants displayed only a modest (less than 5-fold) increase in resistance to 1249A8 compared Δ3a7b-Nluc (**Figures 3B and 3C**). Specifically, the NT_50_ values for ARM-B7, ARM-A8, and ARM-B7+A8 against 1249A8 were 5.51 µg/mL, 9.55 µg/mL, and 4.20 µg/mL, respectively, values that are comparable to 2.09 µg/mL for the original Δ3a7b-Nluc.

**Figure 3.**
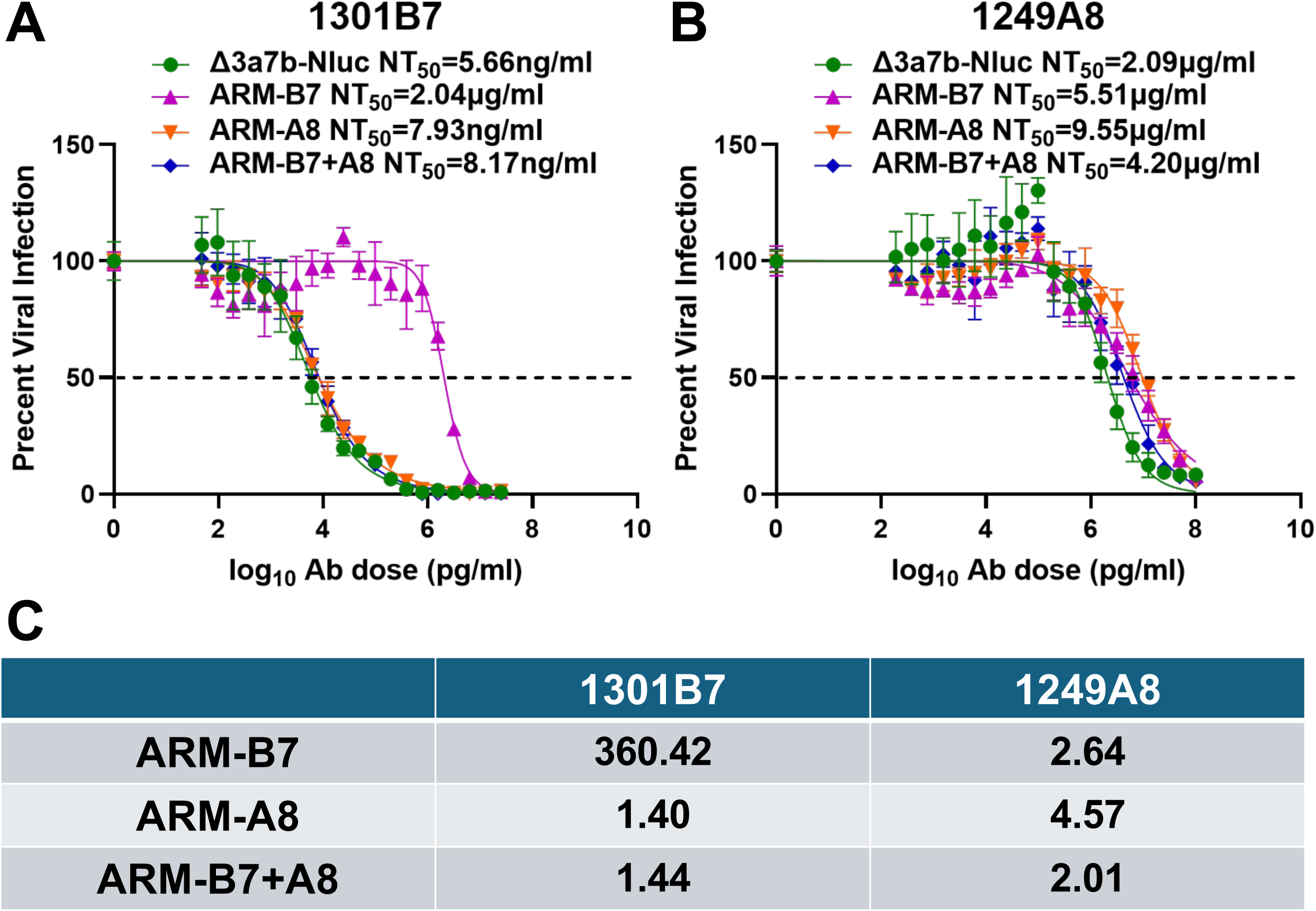
Neutralizing activity of 1301B7 and 1249A8 with parental and ARMs. Plaque reduction neutralization tests (PRNT) were used to assess the sensitivity of the parental virus (Δ3a7b-Nluc) and ARM variants (ARM-B7, ARM-A8, ARM-B7+A8) to NAbs 1301B7 (A) and 1249A8 (B). Neutralization curves were fitted using nonlinear regression (GraphPad Prism) to calculate the half-maximal neutralization titer (NT_50_). Data are shown as mean ± SD. The dashed line indicates 50% inhibition. (C) Fold-change in NT_50_ for each ARMs relative to the parental virus against 1301B7 or 1249A8 NAbs.

### Identification of amino acid mutations responsible for ARMs

To identify the genetic determinants of viral resistance, we conducted next-generation sequencing (NGS) on viral RNA extracted from infected Vero-AT cells, focusing on variants with a frequency >30% (**Figure 4A**). ARM-B7 (selected with 1301B7) contained three mutations in NSP1 (D139Y), NSP13 (V45A), and S (S371F). ARM-A8 (selected with 1249A8) harbored also three mutations in NSP3 (T847I), NSP16 (A34V), and S (T299I). ARM-B7+A8 mutant (selected with the 1301B7 and 1249A8 cocktail) contained four mutations in NSP1 (D139Y), NSP10 (C41W), NSP13 (V45A), and NSP14 (M58I). No mutations were identified in the S glycoprotein of ARM-B7+A8 after serial passages.

**Figure 4.**
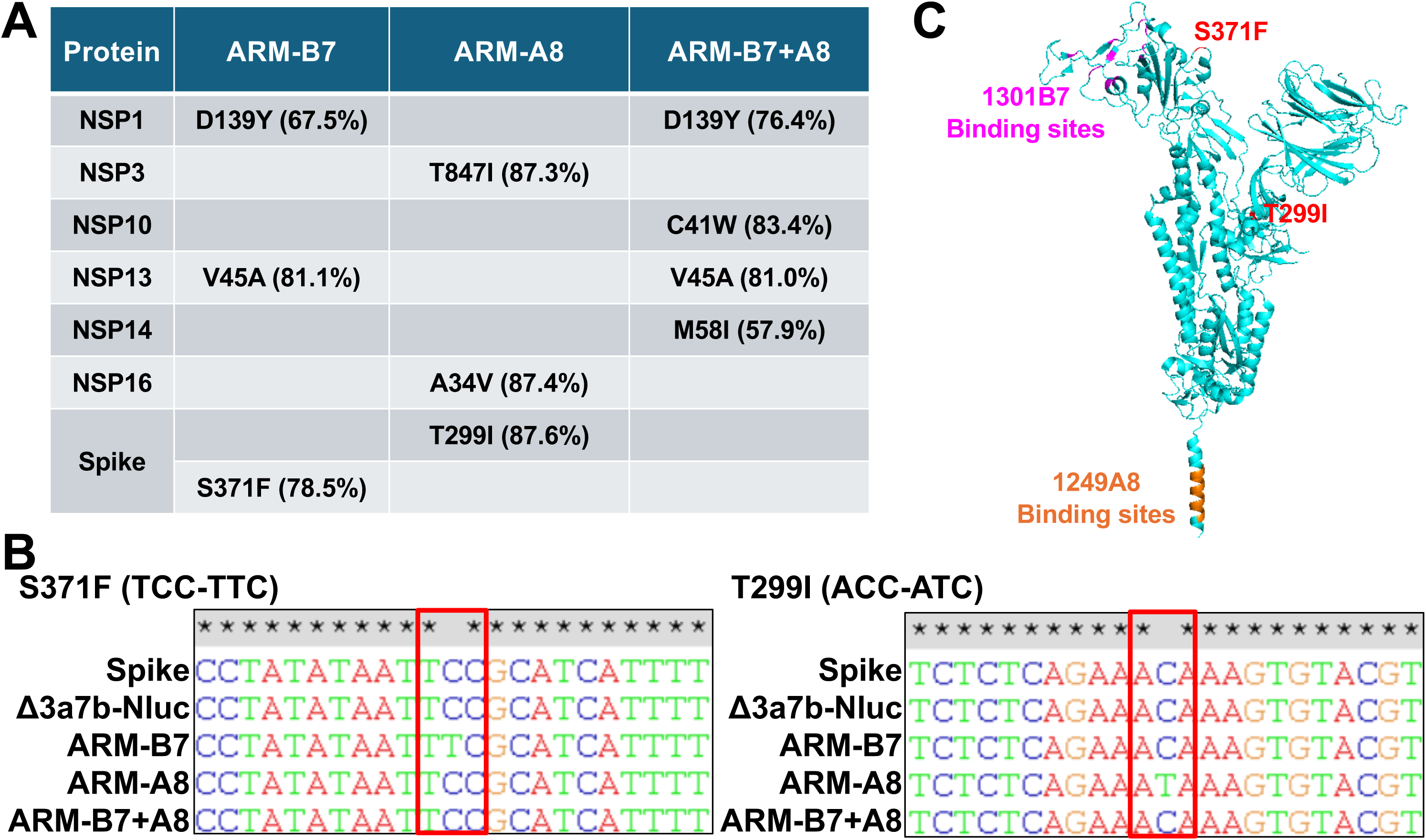
Identification of resistance-conferring mutations in the ARM variants. (A) Amino acid substitutions present at >30% frequency in serially passaged Δ3a7b-Nluc under selective pressure of 1301B7 (ARM-B7), 1249A8 (ARM-A8), or their cocktail (ARM-B7+A8), as identified by NGS. S371F in ARM-B7 and T299I in ARM-A8 within the S protein identified as candidates for conferring resistance. (B) Sanger sequencing chromatograms confirming the presence of the S371F and T299I mutations in the S gene. (C) Structural localization of S371F and T299I mutations (red) within the SARS-CoV-2 S trimer (cyan, PDBID: 6XR8). The binding sites of 1301B7 (magenta) and 1249A8 (orange) are shown.

Since both NAbs target the S protein, we hypothesized that the mutations S371F in ARM-B7 and T299I in ARM-A8 were primarily responsible for the increased neutralizing resistance observed. To further validate our initial NGS results, we conducted Sanger sequencing from RT-PCR products and confirmed the presence of these mutations in the S protein of the respective ARMs (**Figure 4B**). Structural mapping revealed that S371F in ARM-B7 lies outside the 1301B7 binding sites (**Figure 4C**), suggesting an allosteric mechanism of escape rather than a direct disruption of NAb binding. The T299I mutation in ARM-A8, also located outside the 1249A8 epitope, but conferred only minimal resistance to 1249A8 (less than 5-fold).

### S371F mutation confers broad-spectrum resistance across RBD epitope classes

We further investigated the functional impact of S371F mutation by testing its effect on a panel of NAbs targeting distinct epitopes on the S1 RBD of SARS-CoV-2 S protein (**Figure 5**). Compared to the parental Δ3a7b-Nluc (NT_50_=2.84 ng/mL), ARM-B7 mutant showed a greater than 300-fold increase in resistance to Casirivimab (NT_50_ = 872.97 ng/mL) (**Figures 5A and 5D**), which targets a Class I epitope. Resistance was even more pronounced against NAb SC27, which targets a cross-reactive Class I and IV epitope, with ARM-B7 (NT_50_ >10 µg/mL) showing greater than 679-fold reduction in susceptibility (**Figures 5B and 5D**). Furthermore, ARM-B7 mutant also exhibited high-level resistance (NT_50_ >100 µg/mL, >307-fold increase) to Sotrovimab, a Class III-targeting antibody (**Figures 5C and 5D**). This broad, pan-resistance profile indicates that S371F mutation does not merely affect a single epitope but most likely induces a conformational change that compromises the neutralization capacity of antibodies across multiple RBD classes.

**Figure 5.**
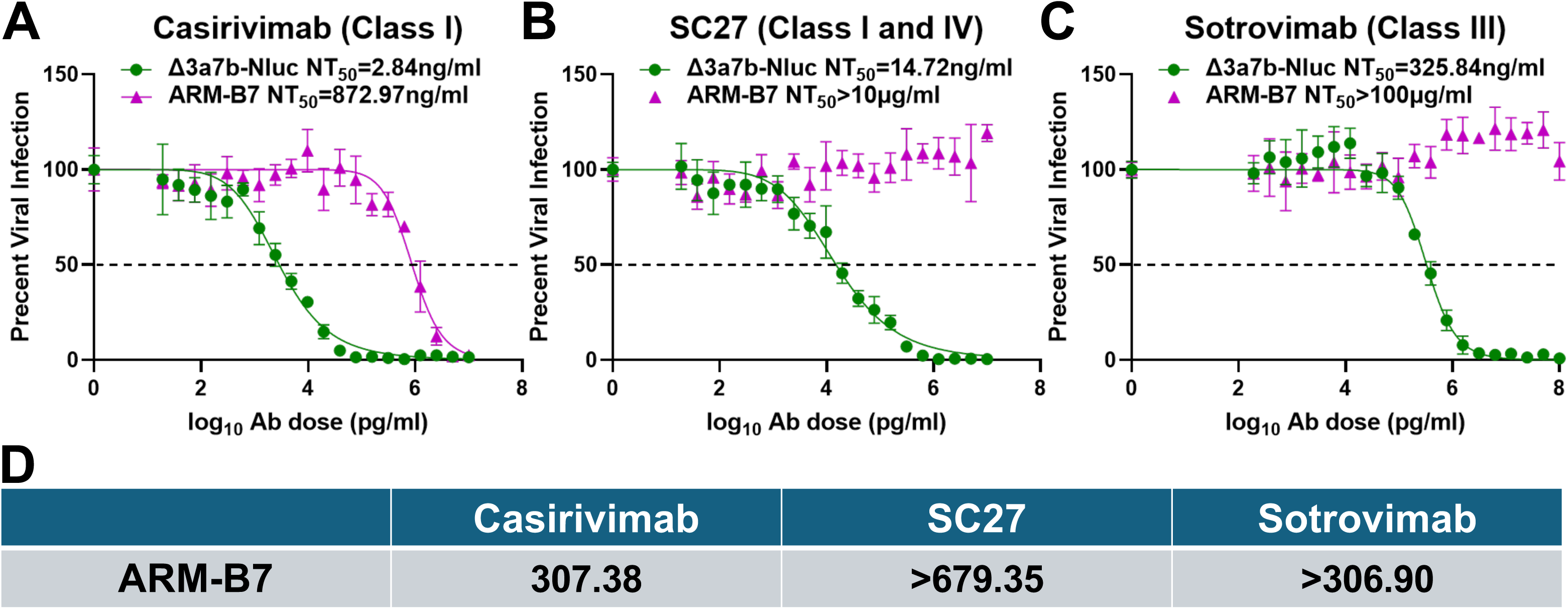
S371F mutation in ARM-B7 confers broad-spectrum resistance to S1 RBD-targeting NAbs. Neutralization sensitivity of the parental Δ3a7b-Nluc virus and the ARM-B7 escape variant to Casirivimab (A), SC27 (B), and Sotrovimab (C) were assessed by PRNT in Vero-AT cells. Dose-response curves were fitted by nonlinear regression to calculate NT_50_ values. Data are shown as mean ± SD. The dashed line indicates 50% inhibition. (D) Fold-change in NT_50_ of ARM-B7 relative to the parental virus against Casirivimab, SC27, and Sotrovimab NAbs.

## DISCUSSION

Therapeutic NAbs have represented a critical intervention for the treatment of SARS-CoV-2 infection, yet all FDA-approved NAb therapies have ultimately been rendered ineffective due to viral evolution (11–13). Although S1 RBD-targeting NAb combination therapies demonstrate reduced susceptibility to escape compared to single NAb treatment, they remain vulnerable to the selection of ARMs even when targeting non-overlapping epitopes due to the S1 RBD’s inherent mutational malleability (29–32). These potent neutralizing capacity of FDA-approved S1 RBD-targeted clinical NAb failure have forced a strategic reassessment of targeted therapeutic options. Our findings provide compelling evidence that NAbs targeting the conserved S2 SH in SARS-CoV-2 S protein exhibit superior resistance profiles for the selection of ARMs, and that combining S2 SH- and S1 RBD-targeting NAbs creates synergistic pressure for the selection of ARMs, significantly outperforming the use of S1 RBD NAb or RBD NAb-combinations. These results establish a new paradigm for NAb therapeutic development against SARS-CoV-2 that emphasize targeting conserved epitopes alongside strategic multi-domain combinations, with careful consideration of epitope landscapes to maximize genetic barriers to resistance, thereby informing the design of more durable evasion-resistant NAb therapeutics.

Unlike most screened NAb resistant mutations that occur directly at or around epitope binding sites (29, 30, 33), the resistance profile of ARM-B7 suggests a distinct mechanism involving antigenic surfaces remodeling of the S protein. This arises from 1301B7’s ability to target relatively conserved regions of the S1 RBD while tolerating for others site diversity, making escape more difficult (16). The S371F mutation identified in this study with ARM-B7 was confirmed to affect multiple S1 RBD NAbs, including Casirivimab (class I) (34), SC27 (class I and IV) (35), and Sotrovimab (class III) (36). Previous research demonstrate that S371F mutation may shift the S from an “up” to a “down” conformation that reduces exposure of the cellular receptor ACE2 binding sites (Class I/II epitopes) (37–40). Additionally, S371F substitution may disrupt the structural environment of the adjacent N343 glycan, thereby compromising the binding efficacy of Class III NAbs (38). Furthermore, as locates in the class IV epitope, S371F may alter the local conformation of the S371-S373-S375 loop, consequently affecting the binding of Class IV NAbs targeting this region (38). First identified in the BA.2 variant and prevalent in subsequent lineages, S371F exemplifies how single-site changes can reshape SARS-CoV-2 S protein structure to evade all four S1 RBD classes NAbs (38, 41). Overall, S371F mediates immune escape not only through direct epitope disruption but also by inducing structural alterations in S protein that undermine NAb recognition, highlighting the strategic versatility of SARS-CoV-2 in evading humoral immunity.

1249A8 potently neutralizes SARS-CoV-2 infection by targeting a highly conserved SH epitope within the S2 stem domain of the S protein, effectively disrupting the conformational changes required for membrane fusion (22, 23). By binding on this structurally conserved region, which remains invariant across the vast majority of SARS-CoV-2 variants, 1249A8 disrupts the post-fusion S protein’s ability to initiate membrane fusion, thereby blocking viral entry (22). Notably, 1249A8 demonstrates broad and potent cross-neutralization not only against diverse SARS-CoV-2 variants but also across multiple β-coronaviruses, including SARS-CoV and MERS-CoV, underscoring its potential as a pan-β-coronavirus NAb therapeutic candidate (23). Notably, the screening process did not yield any highly potent escape mutants, indicating a high barrier to resistance and supporting the broad therapeutic promise of this NAb.

For our studies, we developed a Nluc-expressing attenuated recombinant SARS-CoV-2 (Δ3a7b-Nluc) incorporating two modifications to facilitate easy identification of antiviral resistance under safe experimental conditions (26). First, the deletion of accessory ORF3a and ORF7b proteins achieves viral attenuation while preserving viral replication. Second, the expression of Nluc enables sensitive and quantitative monitoring of viral propagation and identification of ARMs (25–28). This novel platform effectively addresses the biosafety constraints inherent in conducting similar experiments using WT SARS-CoV-2 while maintaining bona-fide SARS-CoV-2 replication rather than pseudotyped approaches, that are essential to accurately identify authentic viral resistant mutants. Moreover, this approach is compatible with high-throughput analysis for the identification of ARMs, thereby offering a versatile and safer standard for resistance surveillance and therapeutic development that can also be safely used to identify ARMs but individual or libraries of antibodies but also with sera samples from naturally infected or vaccinated individuals.

In summary, this study establishes the utility of Δ3a7b-Nluc as a safe and effective approach for identifying ARMs circumventing the biosafety limitations associated with using WT SARS-CoV-2. Our results reveal that NAbs targeting the conserved S2 SH region present a higher genetic barrier to resistance compared to S1 RBD-directed NAbs, and the rational of designed NAb cocktail therapies targeting both S1 RBD and S2 SH to effectively suppress the emergence of SARS-CoV-2 ARMs commonly observed using S1 RBD-targeting monotherapies or S1 RBD NAbs-combinations. Altogether, these findings provide critical insights for developing next-generation resistance-evading NAb therapeutics and demonstrate the value of using attenuated viral platforms to safe and effective resistance analysis without potential biosafety concerns.

## MATERIALS AND METHODS

### Biosafety

*In vitro* experiments with Δ3a7b-Nluc were performed under Biosafety Level 2+ (BSL-2+) containment laboratories. These studies were reviewed and approved by Texas BioMed’s Institutional Biosafety Committee (IBC).

### Cells, viruses and antibodies

Vero cells stably expressing human angiotensin-converting enzyme 2 (hACE2) and transmembrane protease, serine 2 (TMPRSS2) (Vero-AT cells) were acquired from BEI Resources and cultured in Dulbecco’s modified Eagle’s medium (DMEM) supplemented with 10% fetal bovine serum (FBS; VWR), 100 U/mL penicillin-streptomycin (Corning), and 10 µg/mL puromycin (InvivoGen) to maintain selection pressure. The Δ3a7b-Nluc and 1301B7, 1249A8, and SC27 NAbs were previously described (16, 23, 26, 35). Casirivimab (REGN10933) and sotrovimab (S309) were bought from MedChemExpress.

### Isolation of ARM

Confluent monolayers of Vero-AT cells in 12-well plates (triplicate wells) were infected with Δ3a7b-Nluc at 100–200 plaque forming units (PFU)/well and incubated for 1 hour at 37°C under 5% CO_₂_. Following viral adsorption, cells were washed with PBS and maintained in culture medium containing increasing concentrations of 1301B7 (NT_50_ = 5.66 ng/mL), 1249A8 (NT_50_ = 2.09 µg/mL), or a combination of both NAbs. After 72 hours, supernatants from the highest antibody concentration exhibiting ≥50% cytopathic effect (CPE) were harvested, and viral titers were quantified via Nluc activity to initiate subsequent passages. Following seven serial passages under escalating NAb conditions, P7 Δ3a7b-Nluc antibody-resistant mutants (ARMs) were collected, amplified in Vero-AT cells, and stored at –80°C for further analysis.

### Half-maximal neutralizing antibody titer (NT_50_)

Plaque reduction neutralization tests (PRNT) were performed by incubating 100–200 PFU/well of virus with serially diluted antibodies for 1 hour at 37°C: 1301B7 starting concentration of 25 µg/mL, 1249A8 starting concentration of 100 µg/mL, casirivimab starting concentration of 10 µg/mL, SC27 starting concentration of 10 µg/mL, and sotrovimab starting concentration of 100 µg/mL. Confluent Vero-AT cells in 96-well plates (quadruplicate wells per condition) were then infected with the antibody-virus mixtures and incubated at 37°C under 5% CO_₂_ for 1 hour. Following viral adsorption, the inoculum was replaced with post-infection medium containing 1% Avicel, and cells were further incubated under the same conditions. At 16 hours post-infection, cells were fixed with 10% formalin for 24 hours, washed with PBS, and permeabilized with 0.5% Triton X-100 for 15 minutes at room temperature. After additional washing, cells were immunostained using the SARS-CoV N protein-specific cross-reactive monoclonal antibody 1C7C7 (1 μg/mL), followed by detection with the Vectastain ABC-HRP kit and DAB Substrate Kit (Vector Laboratories) according to manufacturer protocols. Viral plaques were detected and quantified using a ChemiDoc MP Imaging System.

### Plaque assay and immunostaining

Confluent monolayers of Vero-AT cells in 6-well plates were infected with 10-fold serial dilutions of the indicated viruses for 1 hour at 37°C under 5% CO_₂_. Following viral adsorption, cells were overlaid with media containing 1% agar and incubated under identical conditions for 72 hours. Cells were subsequently fixed overnight with 10% formaldehyde solution, permeabilized with 0.5% Triton X-100 for 15 minutes at room temperature and subjected to immunostaining with 1C7C7 (1 μg/mL). Detection was performed with the Vectastain ABC-HRP kit and DAB Substrate Kit (Vector Laboratories) according to manufacturer specifications. Plates were scanned and imaged using a ChemiDoc MP Imaging System. Wells were stained with crystal violet for additional visualization and imaged with a ChemiDoc MP Imaging System. The titer of the virus was determined by the number of viral plaques at the corresponding dilution.

### Sequencing

Viral genome sequences were confirmed by whole genome sequencing using the MinION platform (Oxford Nanopore Technologies). Briefly, total RNA was extracted from infected Vero-AT cells using Trizol reagent (Thermo Fisher Scientific) following the manufacturer’s recommendations. cDNA was generated using the SuperScript IV Vilo Master Mix (Invitrogen). PCR amplification was performed with the Artic V5.3.2. NCOV-2019 Panel (Integrated DNA Technologies) and sample libraries were prepared with the Native Barcoding Kit 24 V14 (SQK-NBD114.24, Oxford NanoporeTechnologies) following the manufacturer’s instructions. Sequencing was performed on R10.4.1 Flow Cells (FLO-MIN114, Oxford Nanopore Technologies) according to the manufacturer’s instructions and reads were analyzed with Geneious Prime software by alignment to the reference sequence. To confirm the presence of the S mutations, the region encoding SARS-CoV-2 S protein was amplified by PCR using the Expand High Fidelity PCR System (Sigma-Aldrich) from cDNA samples. The resulting amplicons were purified and subjected to Sanger sequencing by Plasmidsaurus Inc.

### Statistical analysis

All data represent the means ± standard deviation (SD) for each group and were analyzed with GraphPad Prism.

## ACKNOWLEDGMENTS

We want to thank BEI Resources for providing Vero-AT cells. This work was supported, in part, by the NIH/NIAID R01 AI161363 (M.R.W., J.J.K., and L.M.-S.) and by CRIPT (Center for Research on Influenza Pathogenesis and Transmission), a NIAID-funded Center of Excellence for Influenza Research and Response (CEIRR, contract #75N93021C00014) to A.G.-S.

## FIGURE LEGENDS

**Figure S1. Synergistic neutralizing activity of 1301B7 and 1249A8.** A checkerboard plaque reduction neutralization test (PRNT) was used to evaluate potential synergistic neutralizing activity of 1301B7 and 1249A8 against Δ3a7b-Nluc. The vertical and horizontal axes represent the concentration of 1301B7 and 1249A8 (X-fold NT_50_), respectively. The heatmap represents the plaque count (spots) as a measure of viral infection, where a decrease in spots indicates neutralization.

