## Supplementary figures and images for "A cocktail of SARS-CoV-2 spike stem helix domain and receptor binding domain human monoclonal antibodies prevent the emerge of viral escape mutants"

### Supplemental Figure 1

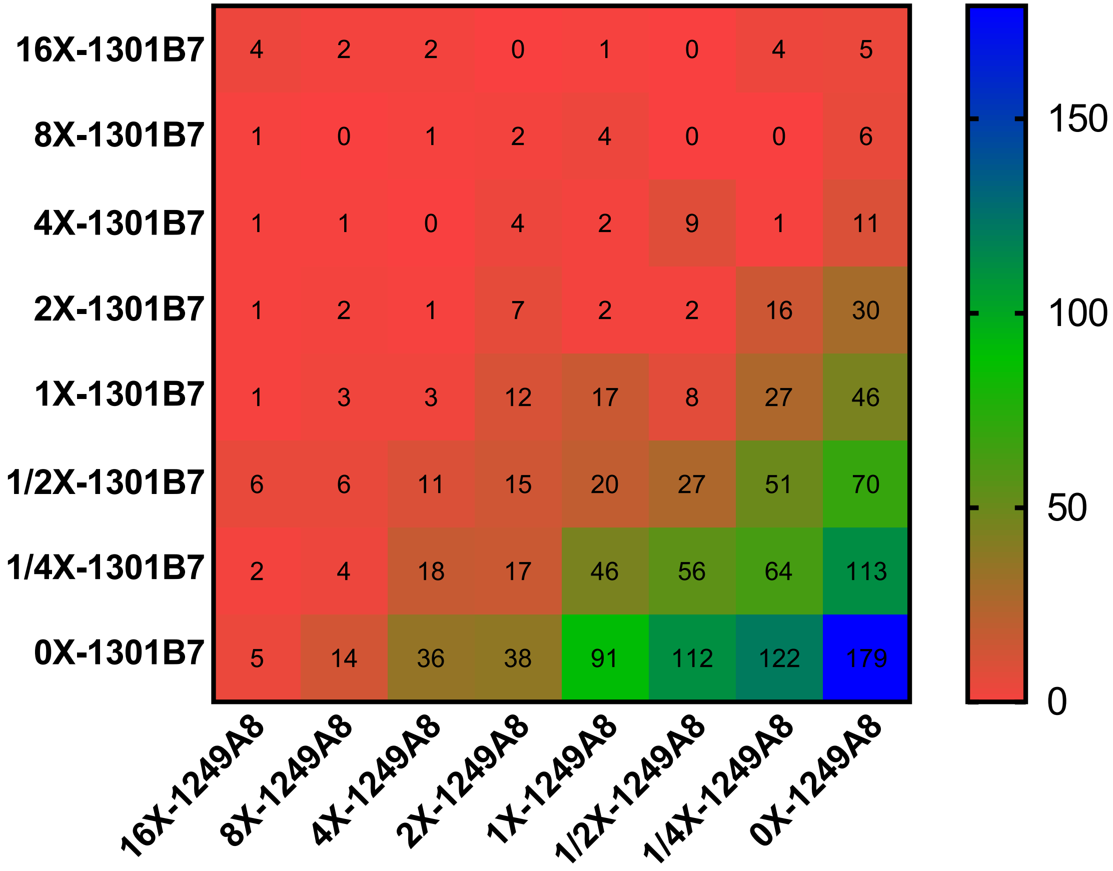
